# Sex- and genotype-effects on nutrient-dependent fitness landscapes in *Drosophila melanogaster*

**DOI:** 10.1101/162107

**Authors:** M. Florencia Camus, Kevin Fowler, Matthew W.D. Piper, Max Reuter

## Abstract

The sexes perform different reproductive roles and have evolved sometimes strikingly different phenotypes. One focal point of adaptive divergence occurs in the context of diet and metabolism, and males and females of a range of species have been shown to require different nutrients to maximise their fitness. Biochemical analyses in *Drosophila melanogaster* have confirmed that dimorphism in dietary requirements is associated with molecular sex-differences in metabolite titres. In addition, they also showed significant within-sex genetic variation in the metabolome. To date however, it is unknown whether this metabolic variation translates into differences in reproductive fitness. The answer to this question is crucial to establish whether genetic variation is selectively neutral or indicative of constraints on sex-specific physiological adaptation and optimisation. Here we assay genetic variation in consumption and metabolic fitness effects by screening male and female fitness of thirty *D. melanogaster* genotypes across four protein-to-carbohydrate ratios. In addition to confirming sexual dimorphism in consumption and fitness, we find significant genetic variation in male and female dietary requirements. Importantly, these differences are not explained by feeding responses and most likely reflect metabolic variation that, in turn, suggest the presence of genetic constraints on metabolic dimorphism.

## 1. Introduction

Males and females perform different reproductive roles and are thus selected for different optimal phenotypes. In response to this divergent selection, the sexes of most species have diverged substantially and show sexual dimorphism in many morphological, molecular and behavioural attributes. One of the key contexts of adaptive divergence between the sexes is diet and metabolism. The composition of the diet has profound effects on lifespan and reproductive output (1, 2) with males and females of many species tailoring their diet to maximise fitness in a sex-specific manner (3). Detailed studies in the fruitfly *Drosophila melanogaster* (4, 5), the field cricket *Teleogryllus commodus* (2, 6), and other insect species (7) have shown that, in order to maximise fitness, females typically require a higher concentration of protein in their diet than males. This nutritional difference between the sexes is consistent with their differing general reproductive roles, where females invest large amounts of resources in the provisioning of eggs but males mainly require energy for the acquisition of mates (4).

In addition to relating their different reproductive roles to nutrition, the sex-specific dietary optima also reflect sex differences in the molecular metabolic machinery. The link between diet and fitness is contingent on many metabolic reactions, as well as on a series of regulatory feedback loops that link the current and anticipated physiological state of individuals to aspects of feeding behaviour and the management of energy stores. Some of these molecular processes have been shown to differ between the sexes. For example, Hoffman et al. (8) characterised the *D. melanogaster* metabolome as a function of fly sex, age and genotype. There was a large effect of sex on metabolite abundance, with 15-20% of the ~1500 assayed metabolites found to differ significantly between males and females. In fact, the real percentage was likely higher as only metabolites that were present in at least 95% of male and female samples were included in the analysis (8). Sex differences in metabolites have also been described in humans (9) with divergence of of almost 80% of the 131 serum metabolites analysed. Moreover, the large majority of these sex differences remained significant after correcting for confounding variables such as age, body mass index, waist-to-hip ratio and lifestyle parameters.

In their study on *D. melanogaster*, Hoffman et al. (8) also detected variation in metabolite concentrations between genotypes, with concentrations of around 10% of metabolites varying significantly between the 15 inbred lines assayed and a similar percentage showing significant age-by-genotype interactions. Genetic variation in metabolites, and diet-induced responses in metabolites have also been found across larvae of different wildtype *D. melanogaster lines* (10, 11). What is currently unknown is whether these genetic effects on the metabolome translate into variation in fitness, and how such fitness effects change with dietary composition. It is conceivable that the differences in titres of at least some metabolites are selectively neutral. This could be the case if the compounds represented intermediate products in metabolic cascades, or if the differences in metabolic fluxes that these measures revealed were usually compensated by behavioural responses that differentially modulated the intake of different nutrients. However, it is also possible that genotypes genuinely vary in the rate and efficiency with which they convert nutrients into reproductive output. The presence of such heritable variation in fitness would indicate that purifying selection on metabolic traits is weak or that genetic polymorphisms in metabolic genes are subject to balancing selection. Either mechanism would prevent metabolism from reaching its adaptive peak and lead to a build-up of a genetic load, where a fraction of the population expresses suboptimal, and hence deleterious, physiologies.

In order to better understand metabolic adaptation and its limits, we need to assess the extent of genetic variation in sex-specific, diet-dependent fitness. In this paper, we build on previous studies of the overall effects of diet on sex-specific fitness (4, 5). We measured male and female diet-dependent fitness of thirty *D. melanogaster* genotypes randomly sampled from the outbred laboratory population LH_M_ (12). In order to assay independent and interactive effects of dietary components on fitness, we used a nutritional geometric framework approach (7) based on a ‘holidic’ diet whose components are completely defined. We estimated genotype-specific male and female fitness surfaces over gradients of dietary protein and carbohydrate ratios
(13, 14) and assessed genetic variation in the parameters that define this surface. We also measured sex- and genotype-specific feeding (the quantity of food consumed) as a function of diet composition, in order to evaluate whether fitness variation arises due to behavioural or physiological responses to diet.

Our results replicate the different sex-specific optima in dietary composition that have been described previously (2, 4). However, we also report significant genetic variation in average male and female dietary requirements, and find contrasting patterns between the male and female requirements of individual genotypes, ranging from overlapping to significantly displaced optima of the sexes.

## 2. Material and Methods

### Fly Stock and Maintenance

We used the experimental base population LH_M_ of *D. melanogaster* for our experiments. This population has been maintained as a large outbred population for over 400 non-overlapping generations, and has been used in previous studies of inter-genomic conflict (15-17). The LH_M_ population is maintained on a strict 14-day regime and with constant densities at both the larval (~175 larvae per vial) and the adult stage (56 vials of 16 male and 16 females each). In line with the regular LH_M_ regime, all base stock flies used in our experiments, were reared at 25°C, under a 12h:12h light:dark photoperiod regime, on cornmeal-molasses-yeast-agar food medium.

We used hemiclonal analysis and sampled thirty haploid genomes, consisting of chromosomes X, II and III (the fourth dot chromosome is ignored), from the population. Hemiclonal haplotypes can be maintained intact and expressed in males and females (15, 18). The hemiclonal flies analysed share the complete genomic haplotype, complemented by chromosomes randomly sourced from the base population. Our experiments thus measure the additive breeding values of the hemiclonal genomes (including those due to epistatic interactions between alleles on the hemiclonal chromosomes), averaged across variable genetic complements, and do not include any non-additive dominance variation (16).

### Synthetic Diet

We used a modified liquid version of the synthetic diet described in Piper et al. (19), that is prepared entirely from synthetic components to enable precise control over nutritional value (see Table S1-S3). Previous studies have used diets based on natural components, typically sugar as the carbon source and live or killed yeast as the protein source (20). Such diets offer only approximate control over their composition, because the yeast-based protein component also contains carbohydrates and is required to provide other essential elements (vitamins, minerals, cholesterol, etc.). As a consequence, phenotypic responses to such diets cannot be straightforwardly interpreted in a carbohydrate-to-protein framework as they are confounded by responses to other dietary components. Our use of a holidic diet completely eliminates these problems.

Four artificial liquid diets were made that varied in the ratio of protein (P, incorporated as individual amino acids) and carbohydrate (C, supplied as sucrose), while all other nutritional components were provided in fixed concentrations. Nutrient ratios used were [P:C] – 1:1, 1:2, 1:4, and 1:16, with the final concentration of each diet being 32.5g/L. This means that the concentration of each dietary component within each diet varies depending on the P:C ratio. These ratios were chosen based on previous work by Jensen et al. (4), who identified these nutritional ratios (or nutritional rails) as the most important in differentiating male and female lifetime reproduction optima.

### Diet Assay and Adult Fitness

Virgin flies were collected within five hours post-eclosion using light CO_2_ anaesthesia. Three flies from each sex/genotype were placed into a vial with a 1% agar and water mixture in order to avoid dehydration with the added benefit that it contains no nutritional value. Flies were kept in these vials overnight before being supplied with a 10μl (females) or 50μl (males) microcapillary tube (ringcaps©, Hirschmann) containing one of the four allocated diets. Capillary tubes were replaced daily, and food consumption for each fly trio was recorded for a total period of four days. Flies were exposed to diet treatments in a controlled temperature room (25°C), 12L:12D light cycle and high relative humidity >80%. The rate of evaporation for all diet treatments was measured by using five vials per diet that contained no flies, placed randomly in the constant temperature chamber. The average evaporation per day was used to correct diet consumption for evaporation. Following four days of feeding under these dietary regimes, flies were assayed for fitness. Male and female fitness experiments were jointly run in 4 identical blocks, with each block comprising all experimental genotypes. Between ten and twelve fly trios were measured for each genotype, yielding a total sample size of 30-36 flies per genotype and diet.

### Male Adult Fitness Assay

Male adult fitness was measured using competitive mating trials (similar to [9]), whereby focal experimental males competed with standard competitor males to mate with females. Following the feeding period described above, a focal trio of virgin males was placed into a new vial, along with three virgin competitor males and six virgin females. The competitor males and the females were of LH_M_ genetic background but homozygous for the recessive *bw*^-^ eye-colour allele. The flies were allowed to interact and lay eggs for a period of 24 hours, after which they were discarded from the vials. Eggs were left to develop for 12 days and the subsequent adult offspring in each vial were counted and scored and assigned to either the focal experimental males (if the progeny had red eyes - wildtype) or the competitor males (if the progeny had brown eyes). During the competitive mating trials, flies were provided with molasses-yeast-agar medium that did not contain live yeast, the main source of food for both males and females (21, 22).

### Female Adult Fitness Assay

Female adult fitness was measured as the number of eggs produced over a fixed period of time. Following the feeding period, trios of virgin females were presented with three males from the LH_M_ stock population, and left to mate/oviposit for 18 hours in vials containing a solid agar medium and *ad libitum* food corresponding to their diet treatment provided via capillary tubes. Following removal of the flies at the end of the oviposition period, the total number of eggs laid were determined by taking pictures of the agar surface and counting eggs using the software QuantiFly (23).

### Statistical Analysis

#### Fitness models

Before statistical analysis, we transformed the fitness data to obtain normally distributed datasets. The female fitness values were transformed by x^2/3^, whereas male fitness values were arcsine transformed. Furthermore, as male and female fitness were measured in different units, we standardised them using Z-transformations.

We used a sequential model building approach (5) with the transformed data across both sexes to assess male and female fitness responses to dietary composition and the degree to which sex-specific responses vary between genotypes. We first analysed sex-specific effects of diet consumption across genotypes, to verify whether we could replicate the results of previous studies (2, 4, 5). We compared a reduced model (Model F1) that describes the fitness response surface with fixed effects for the linear, quadratic and cross-product effects of the consumed diet components with a more complete model (Model F2) that also allows for sex-specific deviations of these effects. In addition, both models account for experimental block effects, modelled as a random effect. The models were specified as:

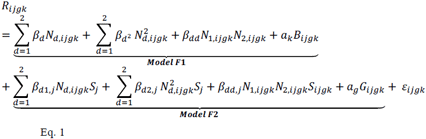

where the underbraces identify terms included in each model. In Equation 1, *R*_*ijgk*_ is the standardised fitness measure of trio *i* of sex *j* and genotype *g* in experimental block *k,N*_*d,ijgk*_ is the amount of dietary component *d* (carbohydrate or protein) consumed by the trio *ijgk* in the feeding period preceding the fitness assay, *β*_d1_ the slope describing how fitness across both sexes changes with consumption of dietary component *d, β*_d2_ is the slope describing how fitness across both sexes changes with the squared consumption of dietary component *d, β*_dd_ is the slope describing how fitness across both sexes changes with the cross-product between dietary components (carbohydrate-by-protein interaction). The sex-specific terms capture deviations [beta_d1j_], [beta_d2j_] and [beta_ddj_] from the general slopes specific to sex *s*_*j*_ of trio *ijk*. B*ijgk* is the value of a categorical variable designating the experimental block of trio *ijgk, a*_*k*_ the value of the coefficient describing the random effect of experimental block (with a~N(0, *σ* _*k*_)), and *ε*_*ijk*_ is the unexplained residual error. Given that our data had been Z-transformed within each sex, we do include neither an intercept nor a term to describe sex differences in mean fitness, as mean fitness is equal to zero overall and in each sex.

In order to assess genetic variation for diet effects on fitness, we added random effect terms to the model in Equation 1 that describe how the flies of different genotypes (hemiclones) vary in their average sex-specific fitness (across all dietary regimes) and linear, quadratic and cross-product effects of carbohydrate and protein intake (Model F3). Again, we built up models in a stepwise manner to a final model

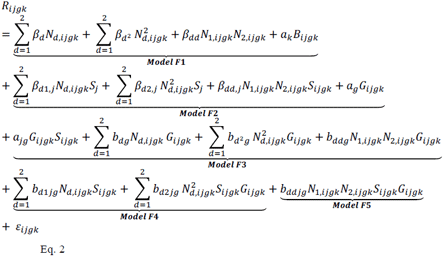

where, as before, the underbraces group terms of specific models. These models include terms describing variation in average sex-specific fitness (where *a*_*g*_ is the effect of the g-th genotype on male and female fitness and *G*_*ijgk*_ designates the genotype identity of trio *igjk*), terms describing genetic variation in the linear parameters of the diet-dependent fitness surface (where *b*_*xg*_ are slopes specific to genotype *g*, with *b*_*xg*_~N(0,*σ*_*xg*_)), and finally, genotype-specific quadratic terms. Models were fitted with maximum likelihood and compared in a pairwise manner (F2 vs. F1, F3 vs. F2, etc.) using parametric bootstrap analysis. We also ran an Analysis of Variance (ANOVA) with type III Sums of Squares using the full model (Eq. 2), in order to assess the significance of individual fixed effect model terms.

In addition to models run on the complete dataset, we also fitted separate models to male and female fitness data. We used these to obtain information on the approximate amounts of fitness variation that can be attributed to the dietary reaction norm of nutritional composition (fixed effects in the mixed-effects models) and to the genotypic variation in dietary responses (random effects in the mixed effects models). To make our approach most straightforward, we fitted fixed effects models including block (as a confounding variable), the scaled quantities of carbohydrate and protein and their interaction (to capture their shared reaction norm), as well as genotype and its interaction with the dietary terms (to capture genotypic effects). We decomposed fitness variance using the (additive) Sums of Squares of these models.

#### Diet consumption models

To examine whether the sexes and/or genotypes varied in the quantity they consumed of each diet, we used a similar model building approach to that used for the fitness data. The basic model (Model C1) expressed diet consumption (*C*_*ifjgk*_-microlitres) of a trio *i* of sex *j* and genotype *g* on diet treatment *f* in block *k* as a function of diet (*D*-fixed effects) and block (*B*-random effect). Model C2 further included a fixed effect for sex (*S*), with Model C3 adding a sex-by-diet interaction as an additional fixed effect to describe how (across genotypes) males and females differ in their average consumption. Further models added random-effect terms describing differences between hemiclones in overall consumption (C4), the effect of diet (C5), the effect of sex (C6) and the interaction of diet and sex (C7)

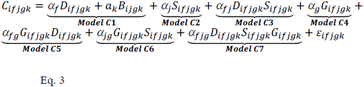

As before, models were fitted with maximum likelihood and compared in a pairwise manner with parametric bootstrap, followed by ANOVA of the full model.

#### Relationship between diet consumption and fitness

We used a permutation approach to determine to what degree fitness variation across genotypes and diets was due to behavioural responses of the genotypes to food (variation in quantity consumed on the different food compositions) or to physiological differences (variation in fitness responses to the same amount of food ingested). Specifically, we permuted—separately for each block, sex and dietary composition—the consumption values across genotypes and then calculated predicted fitness values based on the complete model fitted previously to the fitness data (Model F5). Permutation is valuable in understanding how diet consumption varies with fitness because it will break any associations between behavioural and physiological responses to the different diets. If the variation in fitness is determined by the amount consumed or by a matching of behavioural responses with physiology, then the permutation of consumption data should lead to a lower average predicted fitness and reduced variation in fitness between genotypes. We tested this by generating predicted fitness values for 1000 datasets with permuted consumption data and comparing the distributions of means and variances in fitness across permutations to observed values of these parameters in the original data. P-values were calculated as the proportion of parameter values calculated from the permuted data that equalled or exceeded the values observed in the original dataset. Permutation tests were performed on the entire dataset (males and females), as well as for each sex separately.

All statistical analyses were performed in R version 3.3.2 (24). Mixed models were fitted with the *lmer* function (*lme4* package version 1.1-12, (25)) using maximum likelihood and compared with parametric bootstrap analysis (26) using the *PBmodcomp* function implemented in the package *pbkrtest* (27). ANOVA with type-III Sums of Squares was performed with the *Anova* function from the *car* package (28) following re-fitting of the models with restricted maximum likelihood. We visualised nutritional landscapes based on untransformed data using non-parametric thin-plate splines implemented in the *Fields* (29) package.

## 3. Results

Our study recovered results previously obtained and shows that, averaged across genotypes, males and females differ significantly in their dietary requirements (comparison between Models F1 and F2, P<0.001; detailed inspection of full model: sex × protein × carbohydrate: F = 26.770, p < 0.001, Table 1). Female fitness is maximised by a higher protein intake than male fitness (Figure 1, Table 1) and the parameters describing the shape of the fitness response surface differed significantly between the sexes (Table 2).

**Table 1:**
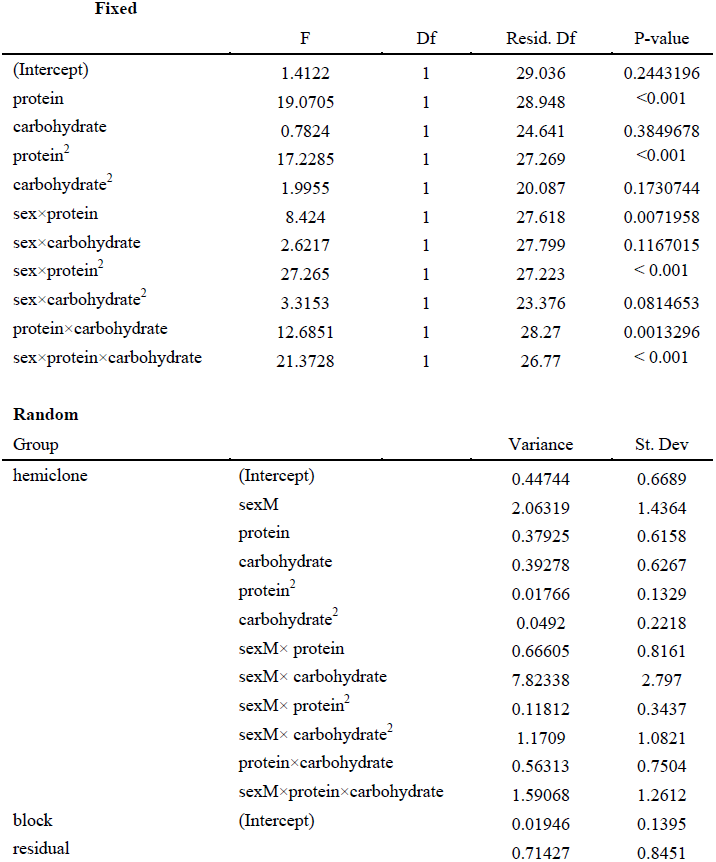
Linear and nonlinear effects of dietary intake and sex on fitness, using the full parametric model (derived from Model F5). The model includes fixed and random effects structure.

**Table 2:** Linear and quadratic and interaction effects of protein (P) and carbohydrate (C) on male and female fitness. Significant effects are represented in bold (P < 0.05).

|  | Linear effects |  | Quadratic Effects |  | Interaction |
| --- | --- | --- | --- | --- | --- |
|  | P | C | P<sup>2</sup> | C<sup>2</sup> | P × C |
| **Male** |  |  |  |  |  |
| Slope ± SE | 0.192 ± 0.246 | -0.866±0.576 | **-0.461±0.103** | 0.383± 0.281 | **0.761±0.260** |
| t | 0.78 | -1.504 | **-4.443** | 1.361 | **2.922** |
| **Female** |  |  |  |  |  |
| Slope ± SE | **0.912±0.194** | 0.437±0.0.438 | **0.3740±0.0651** | 0.1314±0.220 | **-1.016±0.226** |
| t | **4.614** | 0.997 | **5.738** | 0.596 | **-4.479** |

The sign of the linear gradient describes the direction of the relationship between nutrient intake and the response variable (fitness). The quadratic effects capturing concave relationships (troughs in nutritional landscape, slope positive), and convex relationships (peaks on the landscape, slope negative). The sign of the interaction terms indicates changes in the effect of one nutrient with the level of the other nutrient.

**Figure 1:**
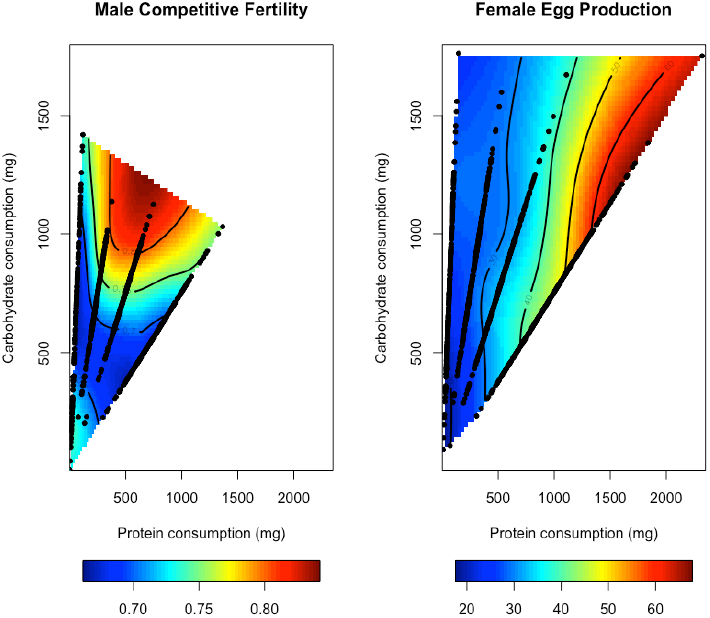
Nutritional landscapes illustrating the effects of protein and carbohydrate intake on the expression of male and female fitness traits. High fitness values are represented by red and low fitness values represented by blue colours. Black dots are individual data points of consumption for the given sex.

Additionally, our data also revealed significant genetic variation in the sex-specific responses to diet (Figure 2). Model comparisons showed that this included variation in average genotype-and sex-specific fitness across all diet treatments (comparison between Models F2 and Model F3, P=0.001), genetic variance in the linear terms describing the shape of the fitness surface across diets (comparison between Models F4 and Model F3, P=0.007) and genetic variation in the quadratic terms of the fitness surface (comparison between Models F5 and Model F4, P=0.005).

**Figure 2:**
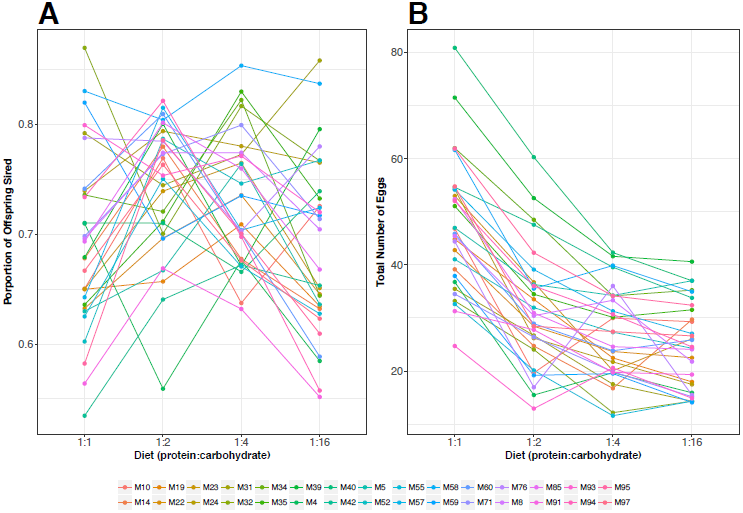
**(A)** Male and **(B)** Female fitness for a suite of 30 genotypes across four different adult diets. Male fitness was measured as male competitive fertility relative to standard competitor males during 24h period after 4d on a dietary regime and female fitness was measured as total number of eggs laid in an 18-hour period post mating (see Methods). Fitness values do not take into account variation in the absolute quantities consumed of protein and carbohydrate (diet), but see Supplementary 2 for genotype-specific fitness landscapes.

Our approximate decomposition of variances based on fixed-effects models suggests that the sexes differ in the contribution of shared (rather than genotype-specific) reaction norms to fitness variation across dietary treatments (males: 4.1%, females: 18.1%). In contrast, the amount of fitness variation that can be attributed to genetic variance in dietary responses is similar in males and females (males: 21.3%, females: 26.2%). These results indicate that overall, the fitness surface across diets is shallower in males than in females, but that the surfaces for males and females of individual genotypes deviate from their sex-specific averages by a similar degree.

Graphical exploration of the fitness surfaces shows that while most genotypes follow largely similar patterns, some genotypes clearly maximise their fitness at very different protein-to-carbohydrate ratios. For example, genotypes M32 (Figure 3), M60 and M94 (Supplementary S2) show males and females having very similar fitness optimum at higher protein levels. On the other hand, some male genotypes required more carbohydrate than the male average to maximise their competitive fitness, resulting in males and females having highly divergent protein-to-carbohydrate optimal ratios (e.g. M31, Figure 3).

**Figure 3:**
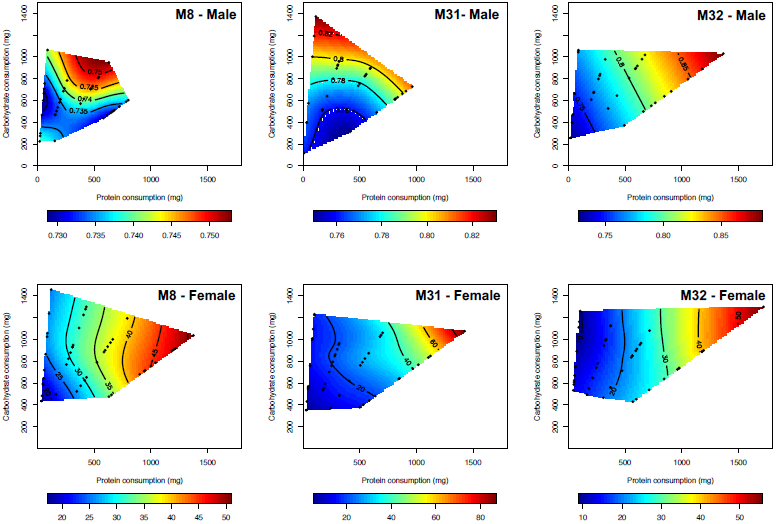
Examples of genotype-specific male and female nutritional fitness landscapes. Hemiclone M8 represents a landscape similar to that found for the population-wide average for males and females (see Figure 1). In contrast, hemiclones M31 and M32 show divergent male optima with similar female optima. M31 males perform best when consuming a diet with a higher carbohydrate concentration, whereas M32 males are most competitive at a higher protein concentration (similar to M32 females). Black dots denote individual data points of consumption for the given sex

For total diet consumption, we find (via model comparison) significant differences between the sexes, with females consuming on average more liquid food than males (comparison between Models C1 and C2, P<0.001, Figure S3-A, Table S3-1). Our results also show differences in consumption between the different diets, with diets containing more protein to carbohydrate being consumed in larger quantities than diets with a higher proportion of carbohydrate (Figure S3-A). Finally, we found high levels of genetic variance for diet consumption within each sex (comparison between Models C6 and C7, P=0.0489, Figure S3-B).

Permutation tests showed that even though genotypes differ in diet-dependent consumption, fitness responses to the dietary treatments was due to physiological, not behavioural, differences between genotypes. Thus, permuting consumption values neither significantly decreased mean predicted fitness nor significantly increased fitness variation, in the entire dataset or when analysing males and females separately (all P > 0.05).

## 4. Discussion

Nutrient acquisition and metabolism are important determinants of fitness components and phenotypic trait expression (3, 5, 6). Our findings shed light on the degree of sex-specific adaptation and optimisation of these processes. By using cytogenetic cloning techniques, we have been able to examine how dietary composition affects male and female fitness of different genotypes of *D. melanogaster*. Our results allow us to assess the overall sexual dimorphism of diet responses and the degree to which genotypes vary in nutritional effects on male and female fitness.

Our results validate previous results showing sex-specific effects of protein and carbohydrate consumption on average sex-specific fitness (4-6). Specifically, male fitness was maximised by a higher proportion of carbohydrates in the diet, while female fitness was highest on a more protein-rich diet. This difference fits with the varying reproductive roles of the sexes. Carbohydrates provide high levels of energy in a short period of time (30) and therefore aid males in obtaining a higher proportion of matings by aggressively pursuing and courting females (31, 32). *Drosophila* females do not suffer from such intense competition as males (33). Instead, reproductive success is mainly determined by the number of viable eggs produced (34), which increases with higher levels of protein (yolk) (35, 36). Similar to previous work, we found that flies altered their feeding behaviour in response to the type of diet provided. Steady state feeding in flies is affected by the interacting forces of the flies’ nutritional history, their mating status and sex, as well as the relative appetitive and satiety values of major dietary macronutrients (37). In our experiments, male and female feeding tended to be higher as the P:C ratio increased, an effect that was also observed in Jensen et al. (4)—one of the few comparable studies to ours because it employed a synthetic (yeast-free) diet at similar P:C ratios and concentrations. When comparing our results with data collected by Jensen et al. (4), it appears that for the concentrations of protein and carbohydrates we used, altered food intake across ratios was principally driven by dietary carbohydrates. This is because increasing carbohydrate content in the food lead to decreasing feeding, irrespective of the P:C ratio (see Figure S3-D for between-study comparison). This could be either because increasing carbohydrates acted as an antifeedant on P:C ratios biased towards higher carbohydrate contents, or because decreasing carbohydrates acted as a phagostimulant on more protein-rich P:C ratios. Distinguishing between these possibilities requires additional behavioural experiments and/or a greater mechanistic understanding of the circuits that drive feeding behaviour.

While identifying known sexual dimorphism in average responses to diet, our experiments also revealed the presence of substantial genetic variation for male and female responses to different diets. Similar to the dimorphism we observe, genetic variation occurs at two levels, in the behavioural responses to diets and the fitness achieved on the selected nutritional ratio. Genetic variation in diet dependent feeding behaviour has been previously described in *D. melanogaster*. Garlapow et al. (38) detected sexual dimorphism for consumption of a single diet for lines from the *Drosophila* Genetics Reference Panel (DGRP). Furthermore, they found significant genetic variation in the mean and variance for consumption, and mapped these traits to genome-wide SNP variation. Our results go beyond this pattern, showing not only variation in consumption of a single diet, but also in how consumption changes when diet composition is altered. In theory, such genetic variation in feeding responses could also entirely explain the heritable fitness variation we detect. For example, a genotype that reduces feeding on certain dietary compositions could suffer nutrient limitation and thus suffer reduced male and/or female fitness. However, this does not seem to be the case in our lines. Permutation analyses showed that fitness effects were not mediated by behavioural responses to food, and individuals of a given genotype and sex did not show lower or higher fitness on a particular diet because they altered the amount ingested. Instead, fitness variation appears to be due to physiological effects, where genotypes differed in the rate at which they were able to convert dietary input into reproductive output.

Irrespectively of their ultimate cause, the extent of diet-mediated fitness variation is surprising. Logically, dietary fitness responses should be subject to strong purifying selection and genetic variation should be eroded rapidly. This raises the question of which evolutionary mechanism could generate such levels of standing genetic variation. Unless the genetic component of dietary fitness responses is very large, it also seems implausible that this level of genetic variation would occur due to additive variation at mutation-selection balance. However, one possibility is that the efficacy of purifying selection is reduced by epistatic interactions. Studies in *Escherichia coli* and *Saccharomyces cerevisiae* have found positive epistasis (where the deleterious effect of double mutations is smaller than the summed deleterious effects of the contributing single mutations) in metabolic genes (39-41). Positive epistasis is particularly prevalent between essential genes, leading He et al. (39) to suggest that this type of interaction should be more important in higher eukaryotes, where a larger proportion of genes are essential.

Alternatively—or in addition—to epistasis, genetic variation in dietary responses could be actively maintained by balancing selection. One potential mechanism for generating balancing selection is temporal or spatial variation in environmental conditions, leading to frequent shifts in adaptive optima (42). Adaptive trade-offs consistent with such a scenario were demonstrated by Sisodia and Singh (43), who investigated the effects of diet on traits related to thermal adaptation in wild-caught *Drosophila ananassae*. The authors found that some macronutrients were beneficial to resistance to heat stress, while others improved cold tolerance (43). Although a plausible mechanism in principle, environmental fluctuations are unlikely to play a role in our study populations. The LH_M_ flies used in the experiments have been maintained under rigorously standardised environmental conditions for more than twenty years. While there have been slight temporal variations in the exact composition of the culture media, these are unlikely to have selected for the large differences in trait response surfaces that we observed.

A further possibility is that genetic variation could be generated and maintained by sexually antagonistic selection on metabolism and physiology, where shared molecular traits are under selection to fulfil opposing demands in males and females. Sexual antagonism is widespread in populations of *Drosophila* (44) including the LH_M_ population studied here (15). In *D. melanogaster*, sexually antagonistic genetic variation has further been shown to exist for diet choice. Experiments using lines from the *Drosophila* Genetics Reference Panel revealed that preferences for particular carbohydrate-to-protein ratios were positively genetically correlated between the sexes (5), while the optimal choice differed between the sexes. In these circumstances, genotypes that express a choice that is optimal in one sex (e.g., a preference for carbohydrate-rich food in males) tend to express a similar but deleterious choice in the other sex (a preference for carbohydrates in females). Similar effects could occur at the metabolic level, where genotypes may vary in the degree to which their metabolism is honed towards the adaptive needs of one or the other sex.

In conclusion, our finding of genetic variance for fitness responses to diet composition suggests that metabolism and physiology are not at their sex-specific adaptive optima. While we have only demonstrated this in the LH_M_ population, many aspects of our data align with results from other sources, such as the variation in feeding responses (38) or metabolite levels (8) that have been identified in the DGRP. This implies that a physiological load, due to the segregation of deleterious metabolic variants, may be common among flies, and potentially in other organisms. Further research is needed to pinpoint the evolutionary mechanisms that allow such variation to accumulate and potentially be actively maintained. Such work will constitute an interesting bridge between evolutionary studies of sex-specific adaptation and functional genetic analyses of nutrient signalling and metabolism.

## Data, Code and Materials

Data from this manuscript will be uploaded to Dryad upon acceptance of this manuscript.

## Authors’ contributions

MFC, KF, MWDP and MR conceived the study and wrote the manuscript. MFC conducted the experiments. MFC and MR analysed the data.

## Acknowledgements

We thank Mark Hill, Filip Ruzicka, Rebecca Finlay and Leonard Pattenden for insightful discussions and help with running experiments.

